# Maintenance of neurotransmitter identity by Hox proteins through a homeostatic mechanism

**DOI:** 10.1101/2022.05.05.490791

**Authors:** Weidong Feng, Honorine Destain, Jayson J Smith, Paschalis Kratsios

**Affiliations:** Department of Neurobiology, University of Chicago, Chicago, IL, USA; University of Chicago Neuroscience Institute, Chicago, IL, USA; Committee on Development, Regeneration, and Stem Cell Biology, University of Chicago, Chicago, IL, USA

## Abstract

Hox transcription factors play fundamental roles during early patterning, but they are also expressed continuously – from embryo through adulthood – in the nervous system. The functional significance of their sustained expression remains unclear. In *C. elegans* motor neurons (MNs), we find that LIN-39 (Scr/Dfd/Hox4-5) is continuously required during post-embryonic life to maintain neurotransmitter identity, a core element of neuronal function. LIN-39 acts directly to co-regulate genes that define cholinergic identity (e.g., *unc-17/VAChT, cho-1/ChT*). We further show that LIN-39, MAB-5 (Antp/Hox6-8) and the transcription factor UNC-3 (Collier/Ebf) operate in a positive feedforward loop to ensure continuous and robust expression of cholinergic identity genes. Finally, we identify a two-component, design principle (Hox transcriptional autoregulation counterbalanced by negative UNC-3 feedback) for homeostatic control of Hox gene expression in adult MNs. These findings uncover a noncanonical role for Hox proteins during post-embryonic life, critically broadening their functional repertoire from early patterning to the control of neurotransmitter identity.

## INTRODUCTION

Information flow in the nervous system critically relies on the ability of distinct neuron types to synthesize and package into synaptic vesicles specific chemical substances, known as neurotransmitters (NTs). Hence, a core functional feature of each neuron type is its NT identity, defined by the co-expression of genes encoding proteins necessary for the synthesis, packaging, and breakdown of a particular NT. Although instances of NT identity switching have been described [1, 2], it is generally accepted that individual neuron types must maintain their NT identity throughout life.

In the case of cholinergic neurons, the enzyme choline acetyltransferase (ChAT) synthesizes acetylcholine (ACh) from its precursor choline, the vesicular ACh transporter (VAChT) packages ACh into synaptic vesicles, the enzyme acetylcholinesterase (AChE) breaks down ACh upon its release into choline, and the choline transporter (ChT) reuptakes choline back into the cholinergic neuron (**Fig. 1A**) [3]. Co-expression of all these proteins throughout the life of a cholinergic neuron (e.g., decades in humans) ensures continuation of ACh biosynthesis, and thereby communication of cholinergic neurons with their post-synaptic targets. Despite ACh being the first NT to be discovered and its widespread use in every animal nervous system [4, 5], how co-expression of ACh pathway proteins is controlled over time, from development through adulthood, is poorly understood. A handful of studies in *C. elegans* and mice identified LIM and POU homeodomain transcription factors as necessary - during development - for ACh pathway gene expression in various types of cholinergic neurons[6-9]. However, the mechanisms that maintain expression of ACh pathway genes during post-embryonic life are largely unknown.

**Figure 1.**
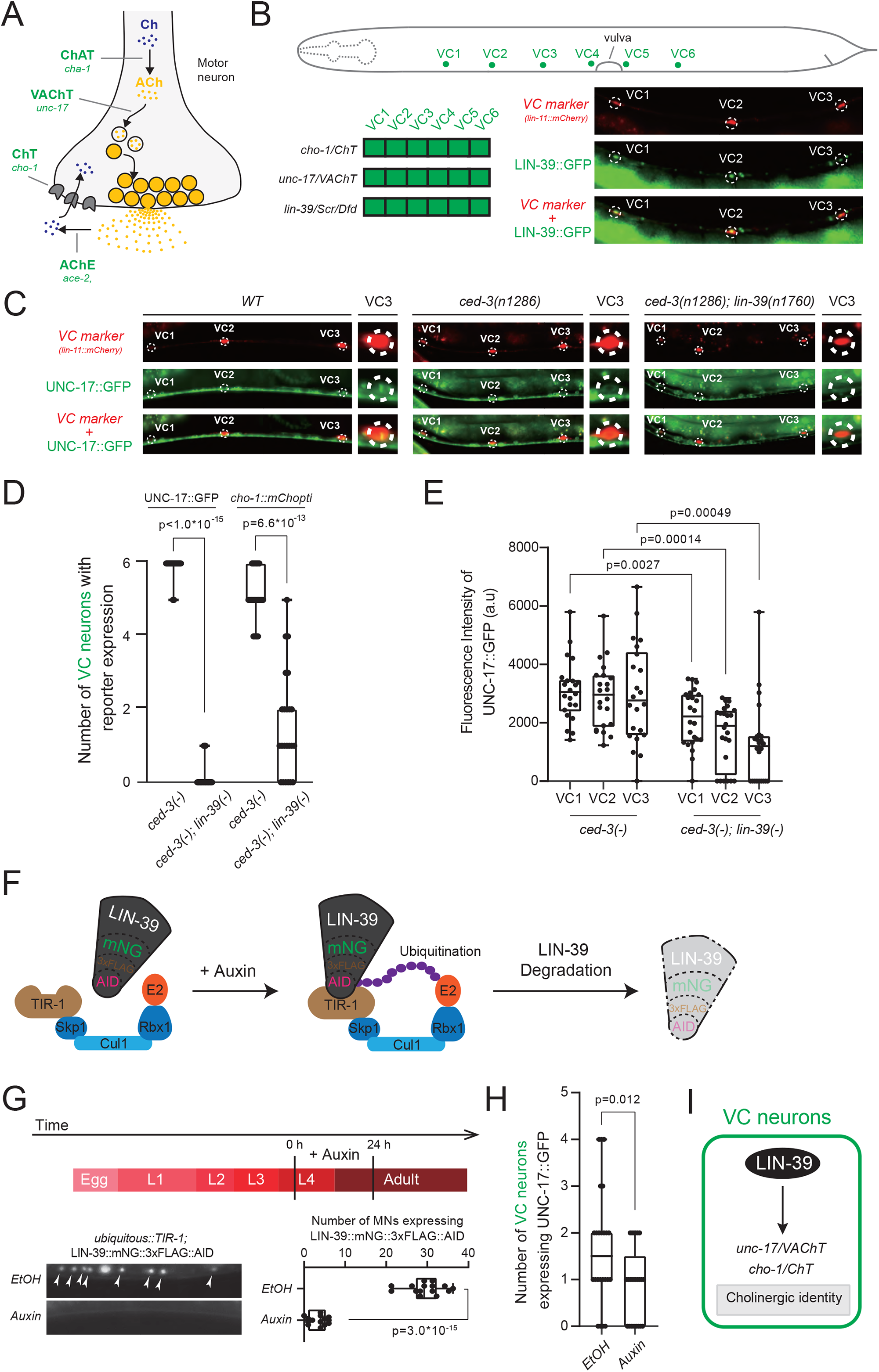
The Hox gene *lin-39 (Scr/Dfd/Hox4-5)* is required to maintain the cholinergic identity of motor neurons necessary for egg laying. **A**. Schematic of a cholinergic motor neuron presynaptic bouton. ACh pathway genes are shown in italics. *cha-1* encodes ChAT: Choline Acetyltransferase. *unc-17* encodes VAChT: Vesicular Acetylcholine Transporter. *cho-1* encodes ChT: Choline Transporter. *ace-2* encodes AChE: Acetylcholinesterase. **B**. *lin-39* is expressed in all VC motor neurons. Top: A cartoon of *C. elegans* hermaphrodite highlighting the cell body of six VC motor neurons in the ventral nerve cord. Bottom left: Colored table demonstrating co-expression of *cho-1, unc-17* and *lin-39* in all VC motor neurons. Bottom right: Representative images of fosmid-based *lin-39* reporter *otIs576* (LIN-39::GFP) in green channel co-expressed with VC specific marker (*lin-11::mCherry*) in red channel with merged channel at the bottom showing VC1-3 expression pattern. Dashed circles indicate cell body positions. **C**. *lin-39* regulates *unc-17* expression in VC motor neurons. Representative images of LIN-39::GFP co-expressed with *lin-11::mCherry* in wild type (*WT*), *ced-3* single mutant (null allele *n1286*), and *ced-3, lin-39* (null allele *n1760*) double mutant background. Red, green and merged channels are shown separately with enlarged portions highlighting expression patterns in VC3. Dashed circles indicate cell body positions. **D**. Quantification of *unc-17* and *cho-1* reporter gene expression under the regulation of *lin-39*. Fosmid-based reporter *otIs544* was used for *cho-1* (as *cho-1::mChopti*). For quantification, box and whisker plots were used with presentation of all data points. Unpaired t-test with Welch’s correction was performed and p-values were annotated. N>20. **E**. Quantification of fluorescence intensity of UNC-17::GFP in VC1-3 in *ced-3* versus *ced-3; lin-39* background. See **Material and Methods** for details on fluorescence intensity quantification. Box and whisker plots were used with presentation of all data points. Unpaired t-test with Welch’s correction was performed and p-values were annotated. N>20. **F**. Schematic of the Auxin induced degradation system. Skp1, Cul1, Rbx1, and E2 are phylogenetically conserved components of the E3 ligase complex. Worms were expressing TIR1 ubiquitously under a ubiquitous promoter (ieSi57). TIR1 is a plant-specific substrate-recognizing subunit of the E3 ligase complex. In the presence of the Auxin, TIR1 binds to the AID fused to LIN-39, leading to ubiquitination and proteasomal degradation of the protein complex of LIN-39::mNG::3xFLAG::AID. **G**. Degradation of LIN-39::mNG::AID upon Auxin treatment. Worms are grown on normal OP50 plates until L4 stages and then transferred to plates containing Auxin or EtOH (as control) and then scored 24 hours later at young adult stages. See **Material and Methods** for more details. Representative images showing LIN-39::mNG::AID expression in ventral nerve cord upon Auxin or EtOH treatment with quantifications were shown below. For quantification, box and whisker plots were used with presentation of all data points. Unpaired t-test with Welch’s correction was performed and p-values were annotated. N>15. **H**. Quantification of the number of VC motor neurons expressing UNC-17::GFP upon Auxin or EtOH treatment (Treatment initiated at L4 stage and images were scored at young adult stage). Box and whisker plots were used with presentation of all data points. Unpaired t-test with Welch’s correction was performed and p-values were annotated. N>20. **I**. Schematic model of Acetylcholine pathway gene regulation in VC motor neurons. LIN-39 regulates *unc-17* and *cho-1* expression determining the cholinergic identity of VC motor neurons.

Motor neurons (MNs) in the spinal cord of vertebrates and ventral nerve cord of many invertebrates use ACh to communicate with their muscle targets. The nematode *C. elegans* contains six different classes (types) of MNs within its ventral nerve cord that are cholinergic [10]. Five classes control locomotion (DA, DB, VA, VB, AS) and are sex-shared (found in *C. elegans* males and hermaphrodites), whereas one class (VC) controls egg-laying and is found only in hermaphrodites. Each MN class contains several members (DA = 9 neurons, DB = 7, VA = 12, VB = 11, AS = 11, VC = 6) that intermingle along the nerve cord. The cholinergic identity of these MNs is defined by the expression of *unc-17* (VAChT), *cha-1* (ChAT), *ace-2* (AChE), and *cho-1* (ChT) (**Fig. 1A**) [12, 13]. The short life cycle of *C. elegans* (∼3 days from embryo to adult), the established methods to inactivate transcription factor activity in the adult, and the availability of reporter animals for all ACh pathway genes offer a unique opportunity to probe *in vivo*, and with single-cell resolution, the molecular mechanisms that maintain cholinergic identity during post-embryonic life.

During early development, members of the HOX family of homeodomain transcription factors control anterior-posterior (A-P) patterning and body plan formation across species[15, 16]. In the nervous system, seminal studies in worms, flies, zebrafish and mice have established critical roles for Hox genes during the early steps of development. Genetic Hox inactivation can affect the specification and/or survival of neuronal progenitors, as well as the differentiation, survival, migration, and/or connectivity of young post-mitotic neurons [17-21]. The focus on early development, however, combined with a lack of temporally controlled Hox inactivation studies that bypass early pleiotropies (e.g., lethality, effects on neural progenitors) has resulted in an incomplete understanding of Hox gene functions in the nervous system. Emerging evidence in *C. elegans, Drosophila*, mice and humans shows that Hox genes are expressed continuously - from development throughout adulthood - in certain neuron types [22-28]. What is the functional significance of sustained Hox gene expression in adult post-mitotic neurons?

Individual neuron types acquire during late development and maintain throughout life their NT identity. Whether and how Hox proteins coordinate the expression of NT pathway genes in developing and/or adult neurons is not known. Here, we show that the *C. elegans* Hox protein LIN-39 (Scr/Dfd/Hox4-5) is continuously required to maintain the expression of ACh pathway genes in nerve cord cholinergic motor neurons, thereby securing their NT identity and function. In motor neurons that control locomotion, LIN-39 cooperates with another Hox protein MAB-5 (Antp/Hox6-8) and the transcription factor UNC-3 (Collier/Ebf) to directly activate the expression of ACh pathway genes. LIN-39 and MAB-5 also regulate the expression levels of *unc-3*, thereby generating a positive feedforward loop that ensures robustness of ACh pathway gene expression throughout life. Furthermore, we propose a homeostatic mechanism that maintains optimal levels of Hox gene expression in motor neurons over time. Because Hox genes are expressed in the developing and adult nervous system of invertebrate and vertebrate animals, the homeostatic mechanism of their regulation and their functional role on maintenance of NT identity may be deeply conserved.

## RESULTS

### The Hox gene *lin-39 (Scr/Dfd/Hox4-5)* is required to maintain the cholinergic identity of motor neurons necessary for egg laying

We initially focused on the hermaphrodite-specific VC motor neurons, for which the regulators of their cholinergic identity, as defined by expression of *unc-17/VAChT* and *cho-1/ChT* (**Suppl. Fig. 1A**) [13], remain unknown. The Hox gene *lin-39* (Scr/Dfd/Hox4-5) is continuously expressed in VC neurons, from the time they are born (larval stage 1, L1) until adulthood (**Fig. 1B**) (**Suppl. Fig. 1B**). However, *lin-39* is required for the survival of VC neurons [29, 30], preventing us from testing its role in the expression cholinergic identity determinants (*unc-17, cho-1)*. To bypass this, we generated double mutant animals for *lin-39* and *ced-3* (because the VC cell death depends on *ced-3* caspase activity)[29, 30], and indeed observed that VC neurons are normally generated in constitutive *lin-39; ced-3* double mutants (**Fig. 1C**). Compared to controls, the number of VC neurons expressing *unc-17* and *cho-1* reporters was significantly reduced in *lin-39 (n1760); ced-3 (n1286)* double mutants in the adult (day 1) stage (**Fig. 1C-D**). Subsequent fluorescent intensity analysis with higher resolution revealed that expression of the cholinergic marker *unc-17::gfp* is significantly reduced, but not completely eliminated (**Fig. 1D**). We note that for the *unc-17* and *cho-1* expression analysis, we used fosmid (∼30kb-long genomic clones)-based reporters, which tend to faithfully recapitulate endogenous gene expression patterns [13]. Altogether, these findings suggest that, during larval development, *lin-39* controls the expression of VC cholinergic identity determinants.

The sustained expression of *lin-39* in VC motor neurons during adulthood prompts the question of whether it is continuously required to maintain cholinergic identity. Because the *lin-39 (n1760)* allele removes gene activity starting in early development, we used an available *lin-39::mNG::3xFLAG::AID* allele that carries the auxin-inducible degron (AID)[31, 32], enabling the depletion of LIN-39 protein in a temporally-controlled manner (**Fig. 1F**). Upon inducing LIN-39 depletion during late larval (L4) and adult stages (**Fig. 1G-H**), we witnessed a statistically significant decrease in the numbers of VC neurons expressing the *unc-17* reporter, indicating *lin-39* is required to maintain at later life stages the expression of *unc-17/VAChT*. Lastly, we examined two additional markers of VC terminal differentiation, *ida-1* (ortholog of human protein tyrosine phosphatase receptor type N [PTPRN]) and *glr-5* (ortholog of human glutamate ionotropic receptor GRIK1). Again, we found that *lin-39* is continuously required to maintain their expression during late larval and adult stages (**Suppl. Fig. 1C**). In conclusion, *lin-39* – in addition to promoting VC survival during L1 – is continuously required to control cholinergic identity and additional features (*ida-1, glr-5*) of VC terminal differentiation (**Fig. 1I**).

### The Hox gene *lin-39 (Scr/Dfd/Hox4-5)* control the cholinergic identity of motor neurons necessary for locomotion

Prompted by our findings in VC motor neurons that control egg-laying, we next asked whether the control of NT identity by *lin-39* extends to motor neurons that control locomotion. Using an endogenous reporter allele (*lin-39::mNG::3xFLAG::AID*), we found *lin-39* to be continuously expressed, from development to adulthood, in 28 of the 39 cholinergic motor neurons necessary for locomotion (**Fig. 2A**) (**Suppl. Fig. 1B**). These neurons survive in animals carrying the same null (strong loss-of-function) *lin-39 (n1760)* allele used in Figure 1, enabling the assessment of putative *lin-39* effects on their cholinergic identity. Indeed, we found a significant decrease in the expression of *cho-1/ChT* (fosmid-based reporter) in these motor neurons (**Fig. 2B-C**). We obtained similar results for two additional cholinergic identity markers using an endogenous reporter for *unc-17 (unc-17::mKate)* and a fosmid-based reporter for *ace-2/AChE* (**Fig. 2D-E**), suggesting *lin-39* co-regulates the expression of several ACh pathway genes in motor neurons necessary for locomotion.

**Figure 2.**
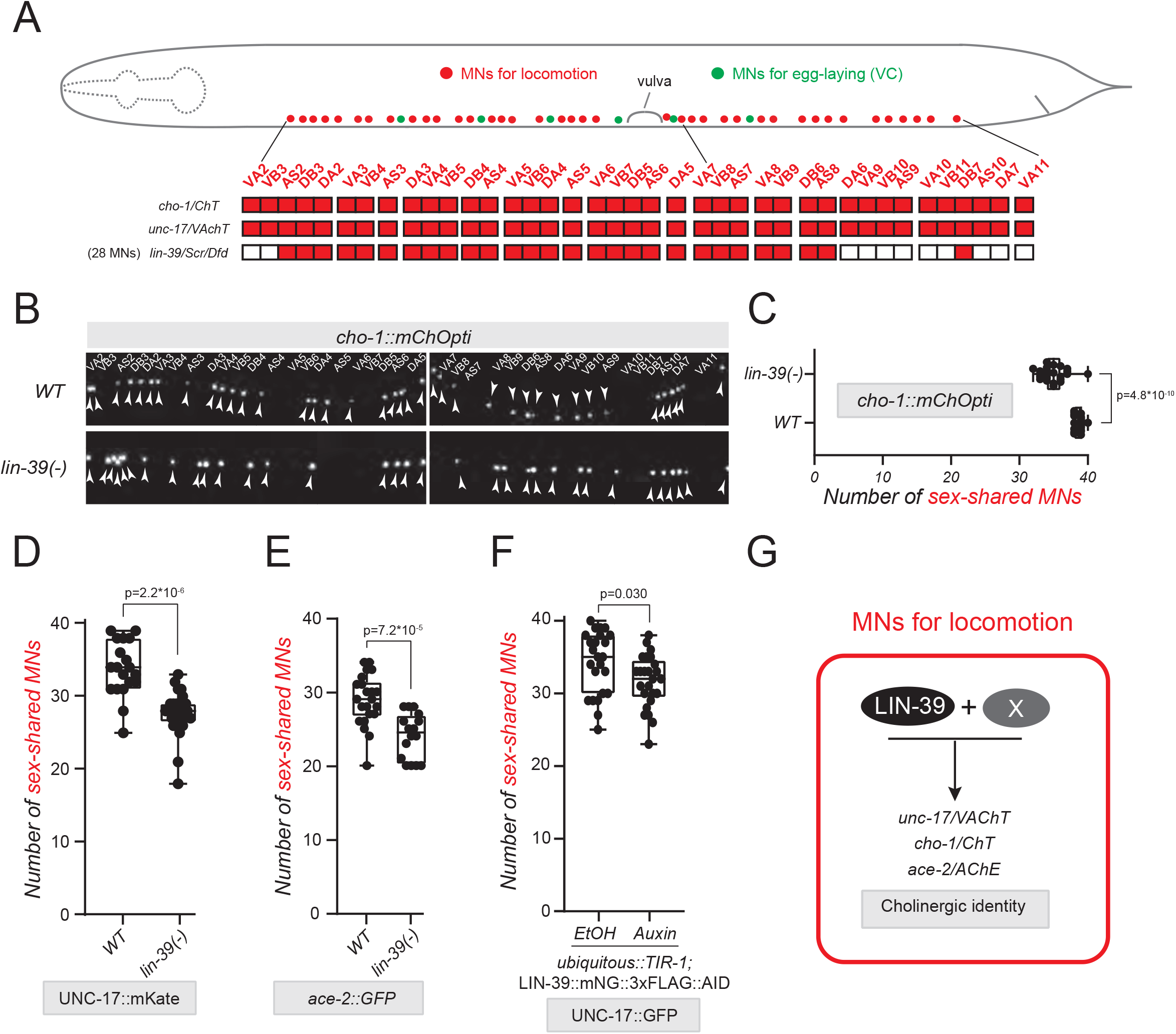
The Hox gene *lin-39 (Scr/Dfd/Hox4-5)* control the cholinergic identity of motor neurons necessary for locomotion. **A**. *lin-39* expression profile in ventral nerve cord sex-shared cholinergic motor neurons with single cell resolution. A cartoon of *C. elegans* hermaphrodite highlighting the cell body of sex-shared cholinergic motor neurons (colored in red) in the ventral nerve cord with cell identity annotated below. Expression profile of *cho-1, unc-17 and lin-39* is illustrated in the color-filled table below. **B**. Representative images of fosmid-based *cho-1* reporter *otIs544 (cho-1::mChOpti)* expression in ventral nerve cord in both *WT* and *lin-39 null* mutant L4 stage hermaphrodite. VC motor neurons are not mature at L4 stage and expression of *cho-1* in VC neurons are invisible. The identity of sex-shared motor neurons is annotated above WT images and cell body positions are pointed out by white arrowheads. Images are separated into anterior/posterior parts (Left/Right) based on vulva positions. **C**. Quantification of the number of motor neurons expressing *cho-1* in panel **B**. Box and whisker plots were used with presentation of all data points. Unpaired t-test with Welch’s correction was performed and p-values were annotated. N>20. **D-E**. Quantification of the number of sex-shared motor neurons expressing *unc-17* (via endogenous reporter *ot907* (UNC-17::mKate) in panel **D**) and *ace-2* (via fosmid-based reporter *otEx4432* (*ace-2::GFP*) in panel **E**) in WT and lin-39 null mutant hermaphrodites at L4 stage. Box and whisker plots were used with presentation of all data points. Unpaired t-test with Welch’s correction was performed and p-values were annotated. N>15. **F**. Quantification of the number of sex-shared motor neurons expressing UNC-17::GFP upon Auxin or EtOH treatment (LIN-39 is endogenously tagged with AID for conditional knock down via Auxin application. Treatment initiated at L4 stage and images were scored at young adult stage). Box and whisker plots were used with presentation of all data points. Unpaired t-test with Welch’s correction was performed and p-values were annotated. N>20. **G**. Schematic model of Acetylcholine pathway gene regulation in sex-shared motor neurons. Partial effects by LIN-39 regulation indicates a role of additional factor X to co-regulate cholinergic identity gene *cho-1, unc-17* and *ace-2*.

Due to its maintenance role in egg-laying motor neurons (**Fig. 1**), we next asked whether *lin-39* is also required to maintain the cholinergic identity of motor neurons critical for locomotion. Again, we found that depletion of LIN-39 specifically during late larval and adult stages resulted in decreased expression of *unc-17/ChAT* (**Fig. 2F**), strongly suggesting that LIN-39 is required to maintain the cholinergic identity of these motor neurons. However, the modest effects observed in the expression of ACh pathway genes indicate that additional factors must co-operate with LIN-39 (**Fig. 2G**).

### The Hox gene *lin-39* cooperates with *mab-5 (Antp, Hox6-8)* and *unc-3* (Collier/Ebf) to control the cholinergic identity of motor neurons

We followed a candidate approach to identify factors that can compensate for the loss of *lin-39* gene activity. First, we reasoned that *mab-5* (*Antp, Hox6-8*), another Hox gene with continuous expression in a subset of *lin-39*-expressing motor neurons (**Suppl. Fig 2**)[25], may work together with *lin-39* to control NT identity features (**Fig. 3A**). Indeed, we observed a greater decrease in the expression of all three cholinergic markers (cho-1/ChT, *unc-17/VAChT, ace-2/AChE*) in double *lin-39(n1760); mab-5 (e1239)* mutants compared to single *lin-39 (n1760)* mutant animals (**Fig. 3B-E**). The residual expression of all these markers in motor neurons of *lin-39; mab-5* double mutants suggests that, besides MAB-5, additional regulatory factors must cooperate with LIN-39 (**Fig. 2B-D**). The transcription factor UNC-3, member of the Collier/Olf/Ebf (COE) family, is selectively expressed in motor neurons that control locomotion (not in egg-laying motor neurons [VC]), and is the only factor previously known to control their cholinergic identity[12]. Hence, if *unc-3* compensates for the combined loss of *lin-39* and *mab-5*, we would predict, in triple mutant *unc-3 (n3435); lin-39 (n1760); mab-5 (e1239)* animals, stronger effects on ACh pathway gene expression compared to single *unc-3* and double *lin-39; mab-5* mutants. Indeed, this was the case for *cho-1/ChT* and *ace-2/AChE* expression (**Fig. 3B-C, E**). Because animals carrying the strong loss-of-function *unc-3* allele *(n3435)* display a dramatic effect on *unc-17/VAChT* expression (UNC-3 is absolutely required for *unc-17/VAChT*), we used a weaker loss-of-function *unc-3* allele *(ot837)[31]*, enabling us to test the notion of cooperation. Indeed, we observed stronger effects on *unc-17/VAChT* expression in *unc-3 (ot837); lin-39 (n1760); mab-5 (e1239)* compared to *unc-3 (ot837)* and double *lin-39; mab-5* mutants (**Fig. 3D**). We extended the analysis of the weaker *unc-3* allele *(ot837)* to *cho-1/ChT* and *ace-2/AChE*, and again observed stronger effects in triple mutant animals (**Fig. 3B-C, E**). Although the magnitude of the observed effects differs in Hox and *unc-3* mutants, our genetic analysis nevertheless suggests that *lin-39, mab-5* and *unc-3* synergize to control the expression of several ACh pathway genes in motor neurons necessary for *C. elegans* locomotion (**Fig. 3F**).

**Figure 3.**
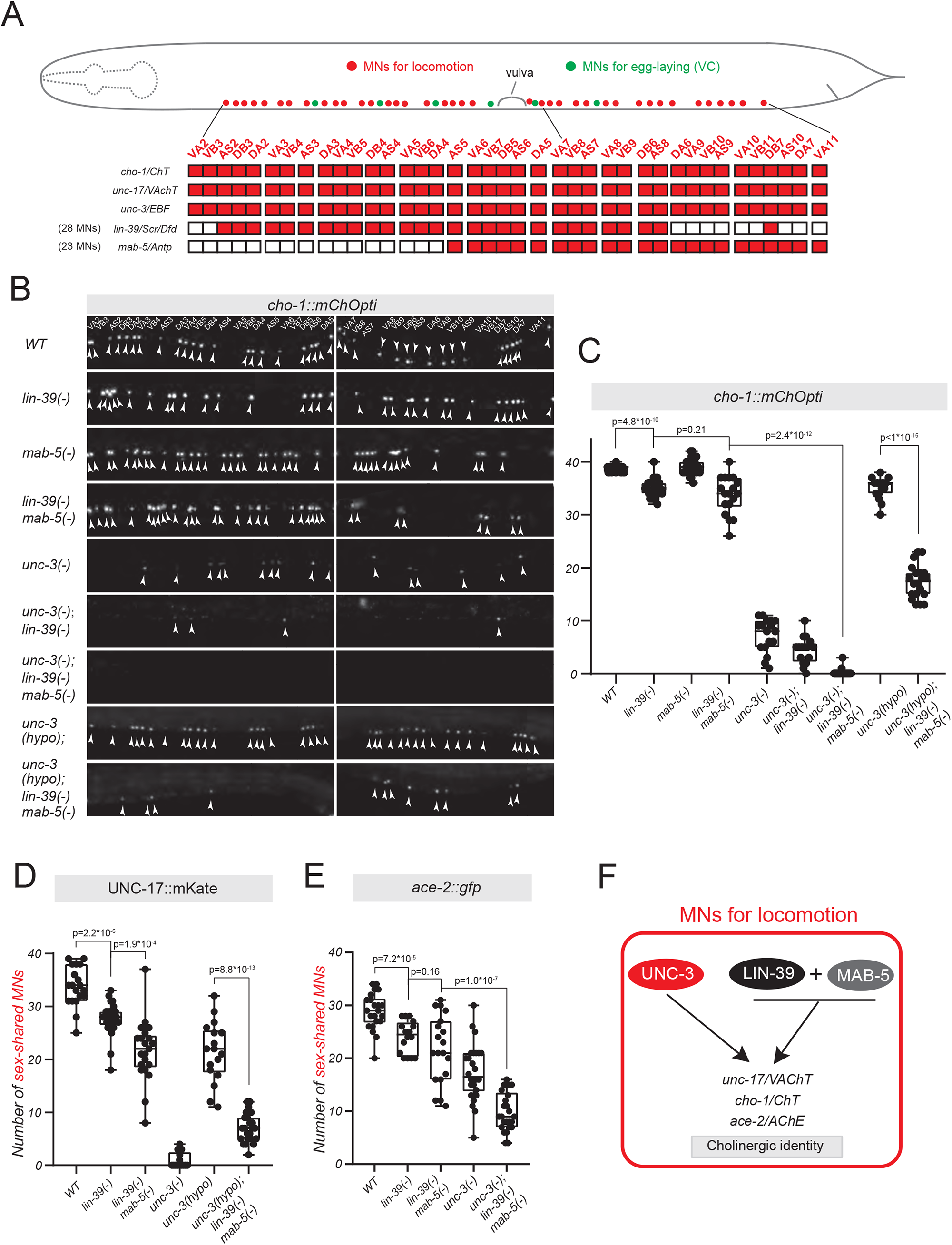
The Hox gene *lin-39* cooperates with *mab-5 (Antp, Hox6-8)* and *unc-3 (Collier/Ebf)* to control the cholinergic identity of motor neurons. **A**. A cartoon of *C. elegans* hermaphrodite highlighting the cell body of sex-shared cholinergic motor neurons (colored in red) in the ventral nerve cord with cell identity annotated below. Expression profile of *cho-1, unc-17, unc-3 and lin-39* is illustrated in the color-filled table below. **B**. Representative images of fosmid-based *cho-1* reporter *otIs544 (cho-1::mChOpti)* expression in L4 stage hermaphrodite ventral nerve cord sex-shared motor neurons in the genetic background of *WT, lin-39(null), mab-5(null), lin-39(null) mab-5(null), unc-3(null), unc-3(null); lin-39(null), unc-3(null); lin-39(null) mab-5(null), unc-3(hypomorph), and unc-3(hypomorph); lin-39(null) mab-5(null). null* is annotated to be *(-)* and *hypomorph* to be *(hypo*). Cell body positions are pointed out by white arrowheads. **C**. Quantification of the number of motor neurons expressing *cho-1* in panel **B**. Box and whisker plots were used with presentation of all data points. Unpaired t-test with Welch’s correction was performed and p-values were annotated. N>15. **D-E**. Quantification of the number of sex-shared motor neurons expressing *unc-17* (via endogenous reporter *ot907* (UNC-17::mKate) in panel **D**) and *ace-2* (via fosmid-based reporter *otEx4432* (*ace-2::GFP*) in panel **E**) in multiple genetic background in hermaphrodites at L4 stage. Box and whisker plots were used with presentation of all data points. Unpaired t-test with Welch’s correction was performed and p-values were annotated. N>15. **F**. Schematic model of Acetylcholine pathway gene regulation in sex-shared motor neurons. *lin-39* cooperates with *mab-5* and *unc-3* to control the cholinergic identity of motor neurons.

### LIN-39 and UNC-3 act through distinct binding sites to activate ACh pathway gene expression in motor neurons

Interrogation of available ChIP-Seq datasets provided biochemical evidence of UNC-3 binding to the *cis*-regulatory regions of ACh pathway genes (*unc-17/VAChT, cho-1/ChT, ace-2/AChE)* (**Fig. 4A**)[33]. These bound regions contain cognate sites for UNC-3, and their previous mutational analysis strongly suggested UNC-3 acts directly to activate the expression of ACh pathway genes in cholinergic motor neurons[12]. Next, we examined ChIP-Seq datasets for LIN-39 and MAB-5 [34], and witnessed largely coincident binding of LIN-39, MAB-5 and UNC-3 to *unc-17/VAChT, cho-1/ChT*, and *ace-2/AChE* loci (**Fig. 4A, D, Suppl. Fig. 3**). We hypothesized that these co-bound regions (putative enhancers) are sufficient to drive reporter (*yfp*) gene expression in motor neurons and found this to be the case. Indeed, a 280bp-long fragment of the *cho-1 cis*-regulatory region, as well as two fragments (1,000bp and 125bp) of the *unc-17* region, are sufficient to drive *yfp* expression in motor neurons (**Fig. 4A-B, D-E**). The *yfp* expression driven by these putative enhancer regions (*cho-1_280bp, unc-17_1,000bp, unc-17_125bp*) is *unc-3*- and Hox-dependent (**Fig. 4B-F**). Similar to our observations with an endogenous reporter for *unc-17* and a fosmid-based reporter for *cho-1* (**Fig. 3C-E**), we corroborated the notion of synergy between UNC-3 and Hox by using the weaker loss-of-function *unc-3* allele *(ot837)*. That is, stronger effects were observed in *unc-3 (ot837); lin-39; mab-5* triple mutants compared to *unc-3 (ot837)* single or *lin-39; mab-5* double mutant animals (**Fig. 4C, E-F**). Altogether, this enhancer analysis in *unc-3* and Hox mutants identifies specific *cis*-regulatory regions that require both UNC-3 and Hox (LIN-39, MAB-5) input to drive of *unc-17/VAChT* and *cho-1/ChT* expression in cholinergic MNs.

**Figure 4.**
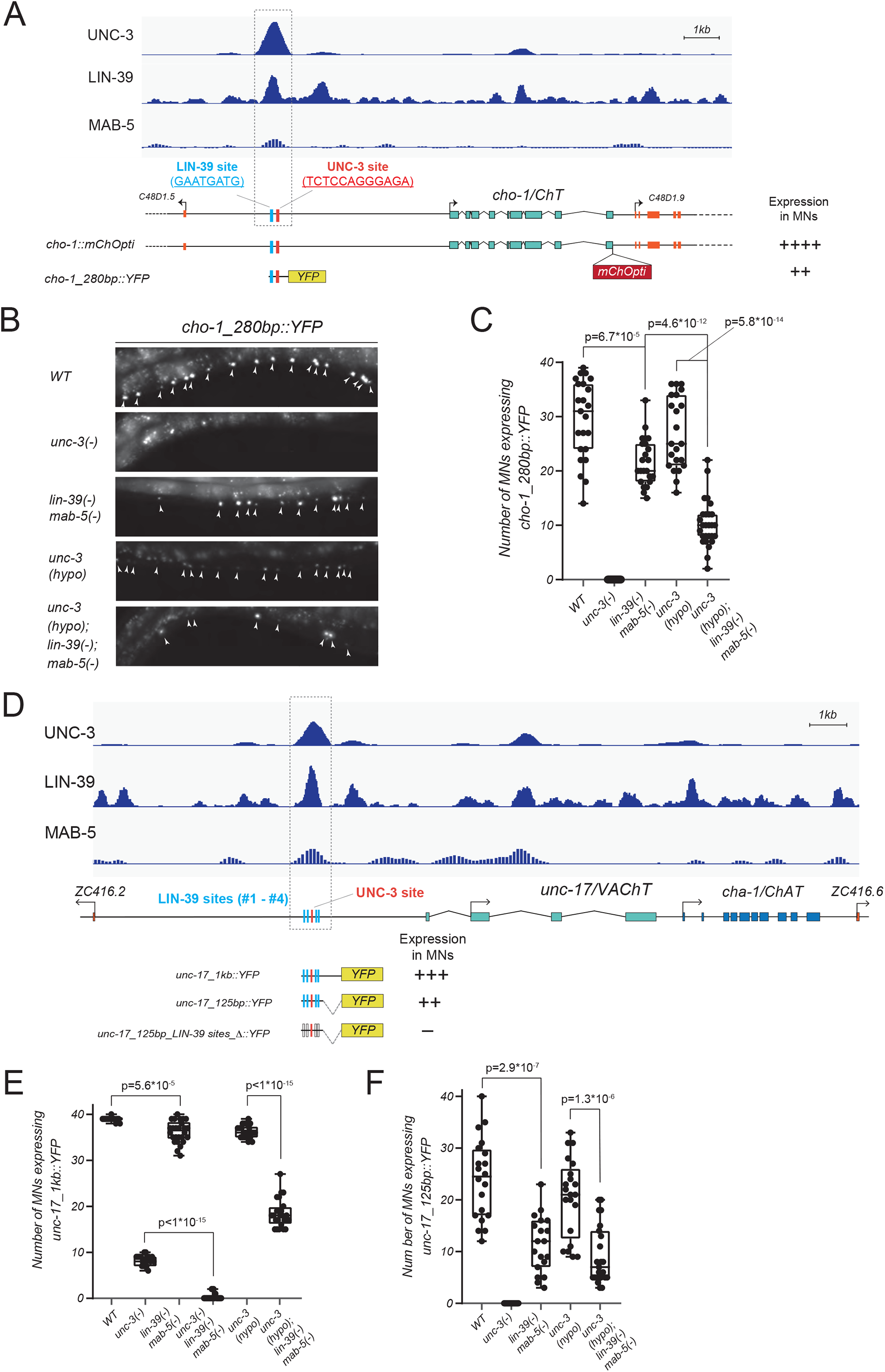
LIN-39 and UNC-3 act through distinct binding sites to activate Acetylcholine pathway gene expression in motor neurons. **A**. Genetic locus of *cho-1* overlay with ChIP-seq tracks of UNC-3, LIN-39 and MAB-5. Molecular nature of *cho-1* reporter gene *cho-1::mChOpti* and *cho-1_280bp::YFP* is annotated overlying below *cho-1* locus. Expression pattern of reporter genes in MNs is indicated with the number of “+”. ++++ indicates expression in more number of MNs with stronger levels; “++” indicates weaker expression in terms of both cell number and expression levels. Within a 280bp cis-regulatory region framed by grey dashed line, a conserved UNC-3 site and a potential LIN-39 site that is predicted through bioinformatics analysis are highlighted in red and blue, respectively. **B**. Representative images of *cho-1* reporter *cho-1_280bp::YFP* expression in L4 stage hermaphrodite ventral nerve cord in the genetic background of *WT, unc-3(null), lin-39(null) mab-5(null), unc-3(hypomorph), and unc-3(hypomorph); lin-39(null) mab-5(null). null* is annotated to be *(-)* and *hypomorph* to be *(hypo*). Cell body positions are pointed out by white arrowheads. **C**. Quantification of the number of motor neurons expressing *cho-1 reporter* in panel **B**. Box and whisker plots were used with presentation of all data points. Unpaired t-test with Welch’s correction was performed and p-values were annotated. N>20. **D**. Genetic locus of *unc-17* overlay with ChIP-seq tracks of UNC-3, LIN-39 and MAB-5. Molecular nature of *unc-17* reporter gene *unc-17_1kb::YFP* and *unc-17_125bp::YFP* as well as its mutagenized version with LIN-39 site deletions is annotated overlying below *unc-17* locus. Expression pattern of reporter genes in MNs is indicated with the number of “+”. “+++” indicates expression in more number of MNs with stronger levels; “++” indicates weaker expression in terms of both cell number and expression levels; “-” indicates completely compromised expression (Data not shown). Within a 125bp cis-regulatory region framed by grey dashed line, a conserved UNC-3 site and 4 potential LIN-39 sites that are predicted through bioinformatics analysis are highlighted in red and blue, respectively. Grey framed LIN-39 sites indicate deletion of sites. **E-F**. Quantification of the number of motor neurons expressing *cho-1* reporter gene *unc-17_1kb::YFP (****E***) and *unc-17_125bp::YFP (****F***) in different genetic background. *null* is annotated to be *(-)* and *hypomorph* to be *(hypo*). Box and whisker plots were used with presentation of all data points. Unpaired t-test with Welch’s correction was performed and p-values were annotated. N>15. **G**. Schematic model of Acetylcholine pathway genes transcriptionally regulated by the synergy of UNC-3, LIN-39 and MAB-5. Red and Blue rectangles indicate the binding of UNC-3 and LIN-39 to gene loci, respectively.

Within the *cho-1_280bp* and *unc-17_125bp* enhancer regions, a single UNC-3 binding site (COE motif) is necessary for reporter gene expression in motor neurons[12]. Next, we bioinformatically searched for the presence of consensus LIN-39 binding sites (GATTGATG), which unlike MAB-5 sites, are well defined in *C. elegans*[34]. We found a single LIN-39 binding site 60bp upstream of the UNC-3 site in the *cho-1_280bp* region (**Fig. 4A**), and four LIN-39 sites flanking the UNC-3 binding site in the *unc-17_125bp* region (**Fig. 4D**). We tested the functional importance of the 4 LIN-39 sites by simultaneously deleting them in the context of transgenic reporter (*unc-17_125bp::yfp)* animals, and found a dramatic decrease in *yfp* expression in motor neurons (**Fig. 4D**). Although this decrease may have occurred due to a change in the physical distancing of *cis*-regulation elements (product of simultaneous deletion of 4 LIN-39 sites) or due to disruption of Hox co-factor binding, our findings nevertheless suggest that LIN-39 and UNC-3 likely act through distinct biding sites. Altogether, this analysis combined with the ChIP-Seq results provide strong evidence that UNC and LIN-39 bind directly to the cis-regulatory region of ACh pathway genes.

### LIN-39 and MAB-5 control *unc-3* expression levels, thereby generating a positive feed-forward loop (FFL) for the control of cholinergic identity

Our findings thus far suggest that Hox proteins LIN-39 and MAB-5, like UNC-3, directly control the expression of ACh pathway genes in motor neurons. However, LIN-39 and MAB-5 also bind extensively on the *cis*-regulatory region of *unc-3* (**Fig. 5A**), raising the possibility of directly regulating its expression. Indeed, expression of an endogenous *gfp* reporter for *unc-3* (UNC-3::GFP protein fusion) is significantly reduced in motor neurons of *lin-39* single mutants, and this effect is exacerbated in *lin-39; mab-5* double mutants (**Fig. 5B-D**). The number of motor neurons that express UNC-3::GFP is lower in *lin-39* and *lin-39; mab-5* animals (**Fig. 5C**). Quantification of UNC-3::GFP fluorescence intensity revealed a striking reduction in motor neurons of Hox mutant animals (34.1% reduction in *lin-39* and 42.5% in *lin-39; mab-5*) (**Fig. 5D**), suggesting *lin-39* and *mab-5* are required for normal levels of UNC-3 expression in motor neurons.

**Figure 5.**
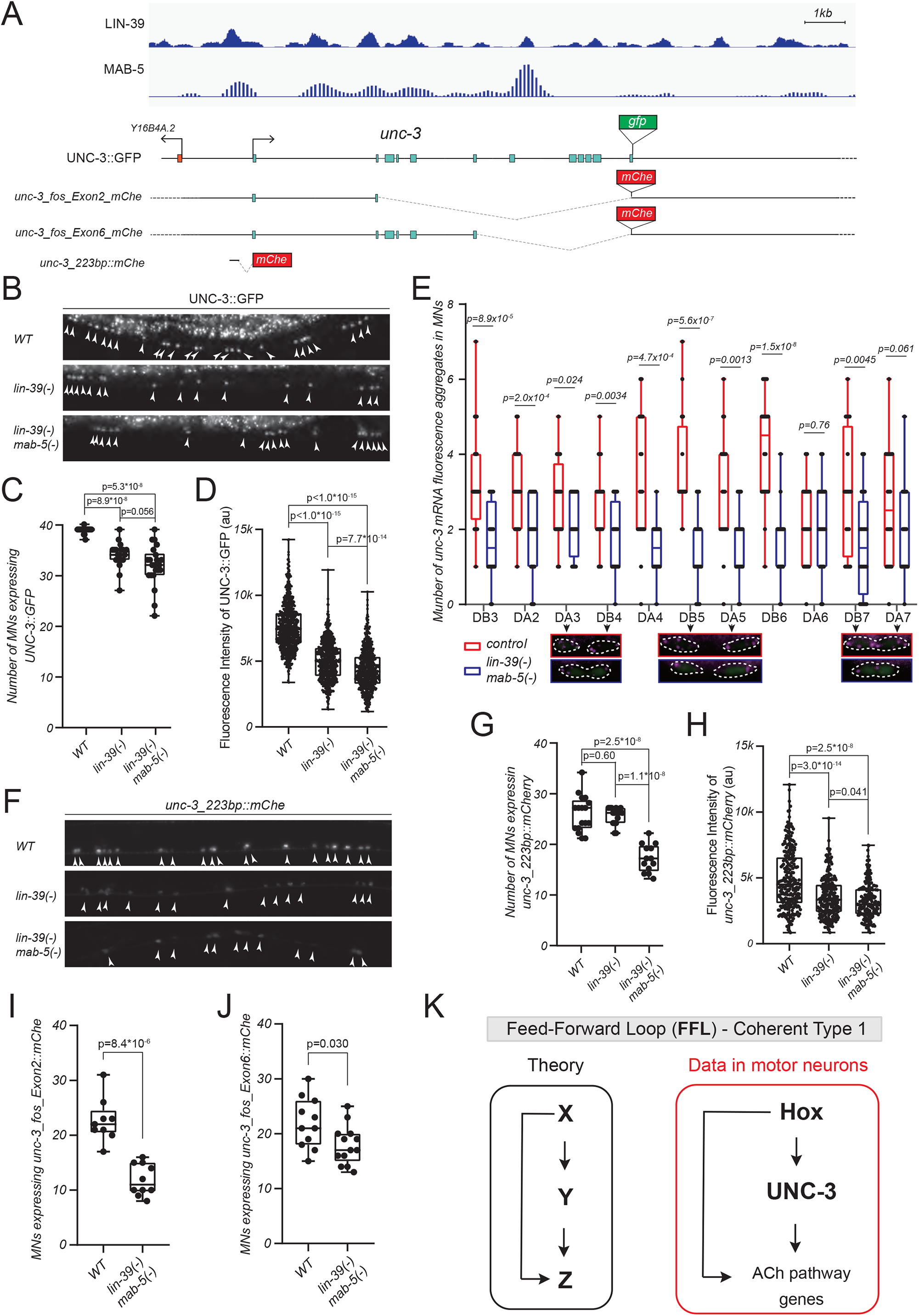
LIN-39 and MAB-5 control *unc-3* expression levels, generating a positive feed-forward loop (FFL) for the control of cholinergic identity. Genetic locus of *unc-3* overlay with ChIP-seq tracks of LIN-39 and MAB-5. Molecular nature of *unc-3* reporter gene UNC-3::GFP, *unc-3_fos_Exon2_mChe, unc-3_fos_Exon6_mChe*, and *unc-3_223bp::mChe* is annotated overlying below *unc-3* locus. **B**. Representative images of UNC-3::GFP expression in L4 stage hermaphrodite ventral nerve cord in the genetic background of *WT, lin-39(null), and lin-39(null) mab-5(null). null* is annotated to be *(-)*. Cell body positions are pointed out by white arrowheads. **C-D**. Quantification of the number of MNs (**C**) or fluorescence intensity (**D**) of UNC-3::GFP in panel Box and whisker plots were used with presentation of all data points. Unpaired t-test with Welch’s correction was performed and p-values were annotated. In panel **C**, N>20. In panel **D**, single neuron expressing UNC-3::GFP in WT or mutant background is scored for fluorescence intensity and plotted as individual data point in graph. N>600. See **Material and Methods** for details on fluorescence intensity quantification. **E**. Quantification of the number of *unc-3* mRNA fluorescence aggregates in individual cholinergic MNs via single molecule RNA Fluorescence in situ hybridization (smRNA-FISH) in WT versus *lin-39(-) mab-5(-)* background with representative zooming-in images of DA3, DB4, DB5, DA5, DB7, and DA7. WT data is colored red versus mutant in blue for quantification. Box and whisker plots were used with presentation of all data points. Unpaired t-test with Welch’s correction was performed and p-values were annotated. N>20. Images are shown in merged channels of *unc-3* mRNA probe stained in magenta with cholinergic MN reporter in green as background. **F**. Representative images of *unc-3_223bp::mChe* expression in L4 stage hermaphrodite ventral nerve cord in the genetic background of *WT, lin-39(-), and lin-39(-) mab-5(-)*. Cell body positions are pointed out by white arrowheads. **G-H**. Quantification of the number of MNs (**G**) or fluorescence intensity (**H**) of *unc-3_223bp::mChe* in panel **F**. Box and whisker plots were used with presentation of all data points. Unpaired t-test with Welch’s correction was performed and p-values were annotated. In panel **G**, N>12. In panel **H**, single neuron expressing *unc-3_223bp::mChe* in WT or mutant background is scored for fluorescence intensity and plotted as individual data point in graph. N>200. See **Material and Methods** for details on fluorescence intensity quantification. **I-J**. Quantification of the number of MNs expressing *unc-3_fos_Exon2_mChe (****I***), and *unc-3_fos_Exon6_mChe (****J****) in WT and lin-39(-) mab-5(-) background*. Box and whisker plots were used with presentation of all data points. Unpaired t-test with Welch’s correction was performed and p-values were annotated. N=10. **K**. Schematic model of a coherent feed-forward loop (FFL) for the control of cholinergic identity by Hox and UNC-3.

The binding of LIN-39 and MAB-5 on the *unc-3* locus strongly suggests these Hox proteins control *unc-3* expression by acting at the level of transcription. Three lines of evidence support this possibility. First, we quantified endogenous *unc-3* mRNA levels with single motor neuron resolution using RNA fluorescent in situ hybridization (RNA FISH)[35]. Compared to wild-type animals, we observed a consistent decrease of *unc-3* mRNA levels in cholinergic motor neurons of *lin-39; mab-5* mutants (**Fig. 5E**). Second, a small 223bp-long *cis*-regulatory element upstream of *unc-3*’s first exon is sufficient to drive reporter gene (*mCherry*) expression in wild-type motor neurons (**Fig. 5A, F**). When we quantified numbers of motor neurons expressing this reporter, as well as its fluorescence intensity, we observed statistically significant differences in Hox mutant animals (**Fig. 5F-H**). Similar to our observation with the UNC-3::GFP fusion (**Fig. 5B-D**), a 27.6% reduction in fluorescence intensity of the *unc-3_223bp::mCherry* reporter is observed in *lin-39* single mutants, and this effect is exacerbated (33.5% reduction) in *lin-39; mab-5* double mutants (**Fig. 5H**). Lastly, we generated two fosmid-based reporters, in which we replaced with *mCherry* two different parts of the *unc-3* locus (2^nd^ intron to last exon, 6^th^ intron to last exon), but left intact the upstream and downstream *cis*-regulatory elements (**Fig. 5A**). In both cases, we found a significant reduction of *mCherry* expression in motor neurons of *lin-39; mab-5* double mutant animals (**Fig. 5I-J**).

In conclusion, Hox proteins (LIN-39, MAB-5) and UNC-3 act directly to activate the expression of ACh pathway genes (*unc-17/VAChT, cho-1/ChT, ace-2/AChE*) in motor neurons. Further, Hox proteins act at the level of transcription to control *unc-3* expression, thereby generating a positive feed-forward loop (FFL) (**Fig. 5K**). In transcriptional regulation networks, the FFL is one of the most widely used network motifs found across species, from yeast to humans [36-38], but its functional significance is experimentally tested primarily in bacteria. In Discussion, we propose that the FFL we discovered in motor neurons is needed to ensure robustness of ACh pathway gene expression.

Similar to the above observations with Hox (LIN-39, MAB-5) and *unc-3*, two other *C. elegans* Hox proteins (CEH-13, EGL-5) control the transcription of another terminal selector (*mec-3*) in peripheral touch receptor neurons [39]. However, our findings on *unc-3* differ in two aspects. First, no evidence was provided that CEH-13 or EGL-5 operate in a FFL in touch neurons. Second, loss of Hox gene (e.g., *ceh-13, egl-5*) activity in touch neurons results in all or none (binary) effects, i.e., 62% of neurons had normal *mec-3* mRNA levels whereas 38% had reduced levels. On the other hand, loss of Hox gene (e.g., *lin-39, mab-5*) activity affected the expression of *unc-3* mRNA and protein in all (100%) motor neurons (**Fig. 5B-E**). Nevertheless, the findings in touch and motor neurons strongly suggest that Hox-mediated transcriptional control of terminal selectors may be a broadly applicable strategy to ensure robustness of gene expression in terminally differentiated neurons.

### Hox genes *lin-39* and *mab-5* maintain their expression in motor neurons through transcriptional autoregulation

To illuminate the gene regulatory mechanisms that maintain, from development through adulthood, Hox gene expression in the nervous system, we initially focused on *lin-39*, which is continuously required to maintain ACh pathway gene (*unc-17/VAChT*) expression in motor neurons (**Fig.1-2**). Available ChIP-Seq data show LIN-39 binding (multiple peaks) at its own locus (**Fig. 6A**), raising the possibility of *lin-39* maintaining its own expression in motor neurons through transcriptional autoregulation. To test this, we first determined which *cis*-regulatory elements of the *lin-39* locus are sufficient to drive reporter (*TagRFP*) expression in motor neurons. Surprisingly, a 6.2kb element upstream of the locus fused to *TagRFP* only shows low levels of expression in a small number of nerve cord motor neurons (∼6) during larval stages (**Fig. 6A**), prompting us to study intronic elements. When intron 1 of *lin-39* was fused to *TagRFP*, we detected robust *TagRFP* expression in ∼23 nerve cord motor neurons. Similar to animals carrying the endogenous *lin-39* reporter allele (*lin-39::mNG::3xFLAG::AID*), these *lin-39* ^*intron 1*^*::TagRFP* animals display continuous expression in both larval and adult motor neurons (**Fig. 6A-B**), thereby enabling us to test the idea of transcriptional autoregulation. Indeed, the number of *TagRFP* expressing motor neurons is significantly reduced in *lin-39* homozygous mutants carrying a null allele (*n1760*) (**Fig. 6B**), suggesting *lin-39* gene activity is necessary for its motor neuron expression.

**Figure 6.**
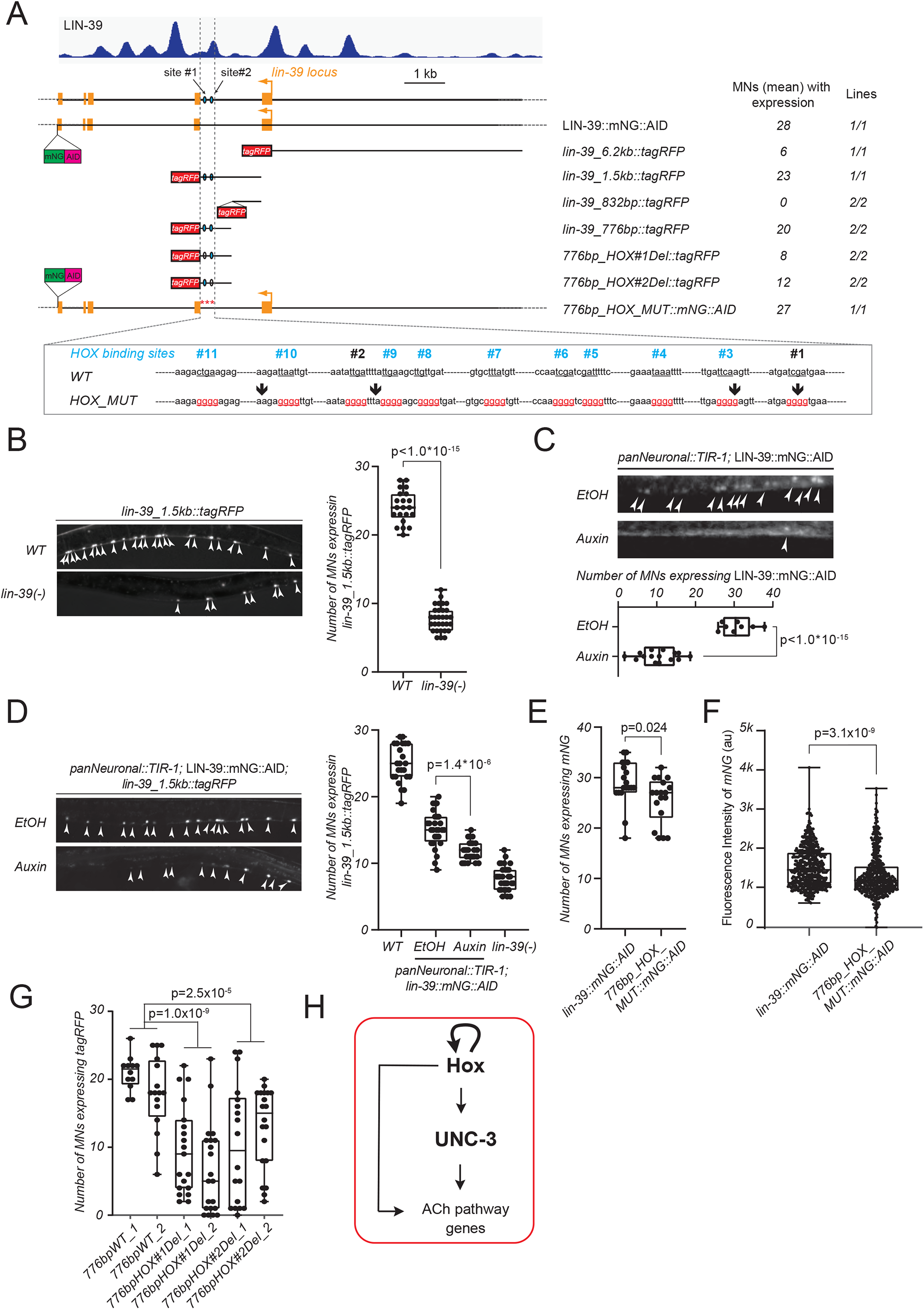
Hox genes *lin-39* and *mab-5* maintain their expression in motor neurons through transcriptional autoregulation. **A**. Genetic locus of *lin-39* overlay with ChIP-seq track of LIN-39. Molecular nature of *lin-39* reporter gene LIN-39::mNG::AID, *lin-39_6*.*2kb::tagRFP, lin-39_1*.*5kb::tagRFP, lin-39_832bp::tagRFP, lin-39_776bp::tagRFP*, mutagenized version (LIN-39 site#1/2 deletion) of *lin-39_776bp::tagRFP* and mutagenized version of LIN-39::mNG::AID as *776bp_HOX_MUT::mNG::AID* is annotated overlying below *lin-39* locus. The mean number of MNs that express the aforementioned reporter genes are indicated on the right side. Details on the mutagenesis in the 776bp cis-regulatory element are highlighted below annotating the 11 LIN-39 binding sites and the mutagenesis scheme verified by Sanger sequencing. **B**. Representative images and corresponding quantification of the number of MNs expressing *lin-39_1*.*5kb::tagRFP* in *WT* versus *lin-39(-)* background. **C**. Degradation of LIN-39::mNG::AID upon Auxin treatment. Worms that express LIN-39::mNG::AID together with TIR protein in all neurons (driven by pan-neuronal promoter *oTti28*) are grown on normal OP50 plates until L2 stage and then transferred to plates containing Auxin or EtOH (as control) and then scored 36 hours later at L4/young adult stages. See **Material and Methods** for more details. Representative images showing LIN-39::mNG::AID expression in ventral nerve cord upon Auxin or EtOH treatment with quantifications were shown below. **D**. Representative images and corresponding quantification of the number of MNs expressing *lin-39_1*.*5kb::tagRFP* treated with EtOH or Auxin as described in panel **C**. For **B-D**, cell body positions are pointed out by white arrowheads. For quantification, box and whisker plots were used with presentation of all data points. Unpaired t-test with Welch’s correction was performed and p-values were annotated. N>20. **E-F**. Quantification of the number of MNs expressing mNG (**E**) or the fluorescence intensity (**F**) of mNG in LIN-39::mNG::AID reporter versus *776bp_HOX_MUT::mNG::AID*, the mutagenized version demonstrated in panel **A**. Box and whisker plots were used with presentation of all data points. Unpaired t-test with Welch’s correction was performed and p-values were annotated. In panel **E**, N>15. In panel **F**, single neuron expressing mNG is scored for fluorescence intensity and plotted as individual data point in graph. N>400. See **Material and Methods** for details on fluorescence intensity quantification. **G**. Quantification of the number of MNs expressing tagRFP in WT 776bp lin-39 reporter versus those with LIN-39 site deletion. Two independent lines are grouped for quantifications. Box and whisker plots were used with presentation of all data points. Unpaired t-test with Welch’s correction was performed and p-values were annotated. N>15.

Because the *lin-39 (n1760)* allele removes gene activity starting in early embryo, the above findings do not address whether LIN-39 is continuously required to maintain its expression at later stages. To test this, we again used the AID system and efficiently depleted LIN-39 protein levels during late larval and adult stages (**Fig. 6C**). Upon auxin administration, we found a significant reduction in the number of motor neurons expressing the *lin-39* ^*intron 1*^*::TagRFP* reporter (**Fig. 6D**), suggesting *lin-39* is continuously required.

These findings pinpoint to the entire intron 1 as a putative *cis*-regulatory region used by LIN-39 to directly control its own expression. Next, we generated transgenic reporter animals that split intron 1 in two fragments, roughly of the same size (832bp and 776bp). Animals carrying the 776bp fragment revealed reporter gene expression in nerve cord motor neurons, but that was not the case for the 832 bp fragment (**Fig. 6A**). Within the 776bp fragment, we found eleven LIN-39 binding sites (motifs) (**Fig. 6A**), and employed CRISPR/Cas9 gene editing to simultaneously mutate all of them in the context of an endogenous reporter for *lin-39* (*lin-39::mNG::3xFLAG::AID*). Combined mutation of all eleven LIN-39 sites led not only to a decrease in the number of motor neurons expressing the endogenous *lin-39::mNG::3xFLAG::AID* reporter (**Fig. 6E**), but also in a reduction of its expression levels (**Fig. 6F**). These findings support a transcriptional autoregulation model where LIN-39 recognizes its cognate binding sites within intron 1 to regulate its expression in nerve cord motor neurons (**Fig. 6H**). Because expression is not completely eliminated in the context of this endogenous reporter with mutated LIN-39 sites, it is likely that other LIN-39 sites and/or additional regulatory factors are involved in the regulation of *lin-39* expression in motor neurons.

To investigate which of the 11 LIN-39 sites are functionally important, we deleted independently the sites with the highest bioinformatic prediction scores. Deletion of either of these two LIN-39 motifs in the context of *lin-39* ^*intron 1b*^*::TagRFP* animals significantly decreased the number of *TagRFP* expressing MNs (**Fig. 6G**), indicating that these motifs are necessary for expression.

Lastly, we asked whether the Hox gene *mab-5* also controls its own expression in motor neurons. Supporting this possibility, ChIP-Seq data show extensive MAB-5 binding at its own locus, and expression of a *mab-5::gfp* reporter is significantly reduced in motor neurons of *mab-5* mutant animals (**Suppl. Fig 4**). Altogether, these findings support the existence of a positive feedback mechanism (transcriptional autoregulation) that operates continuously to maintain Hox gene (e.g., *lin-39*) expression in motor neurons.

### UNC-3 (Collier/Ebf) prevents high levels of Hox gene (*lin-39, mab-5*) expression

If Hox transcriptional autoregulation is perpetuated without negative feedback, high Hox levels will be accumulated in motor neurons with potentially detrimental effects on cell identity and/or function (see Discussion). Hence, there must be negative feedback mechanisms that counterbalance the autoregulatory strategy of *lin-39* and *mab-5*. Because UNC-3 binds to the *lin-39* and *mab-5* loci (**Fig. 7A, Suppl. Fig. 5**), we hypothesized it acts directly to prevent high levels of Hox gene expression in cholinergic motor neurons. Multiple lines of evidence support this idea. First, the expression levels of the endogenous *lin-39* reporter (*lin-39::mNG::3xFLAG::AID*) is increased in motor neurons of *unc-3* null mutant animals (**Fig. 7D**). Second, the *lin-39* ^*intron 1*^*::TagRFP* reporter showed increased *TagRFP* expression in motor neurons of *unc-3* mutants (**Fig. 7E-G**). Lastly, RNA FISH revealed increased levels of *lin-39* mRNA molecules in individual motor neurons of *unc-3* mutants (**Fig. 7H**). Importantly, we extended this analysis to *mab-5*, and obtained similar results – *unc-3* gene activity is necessary to prevent high levels of *mab-5* expression in motor neurons (**Suppl. Fig. 5**).

**Figure 7.**
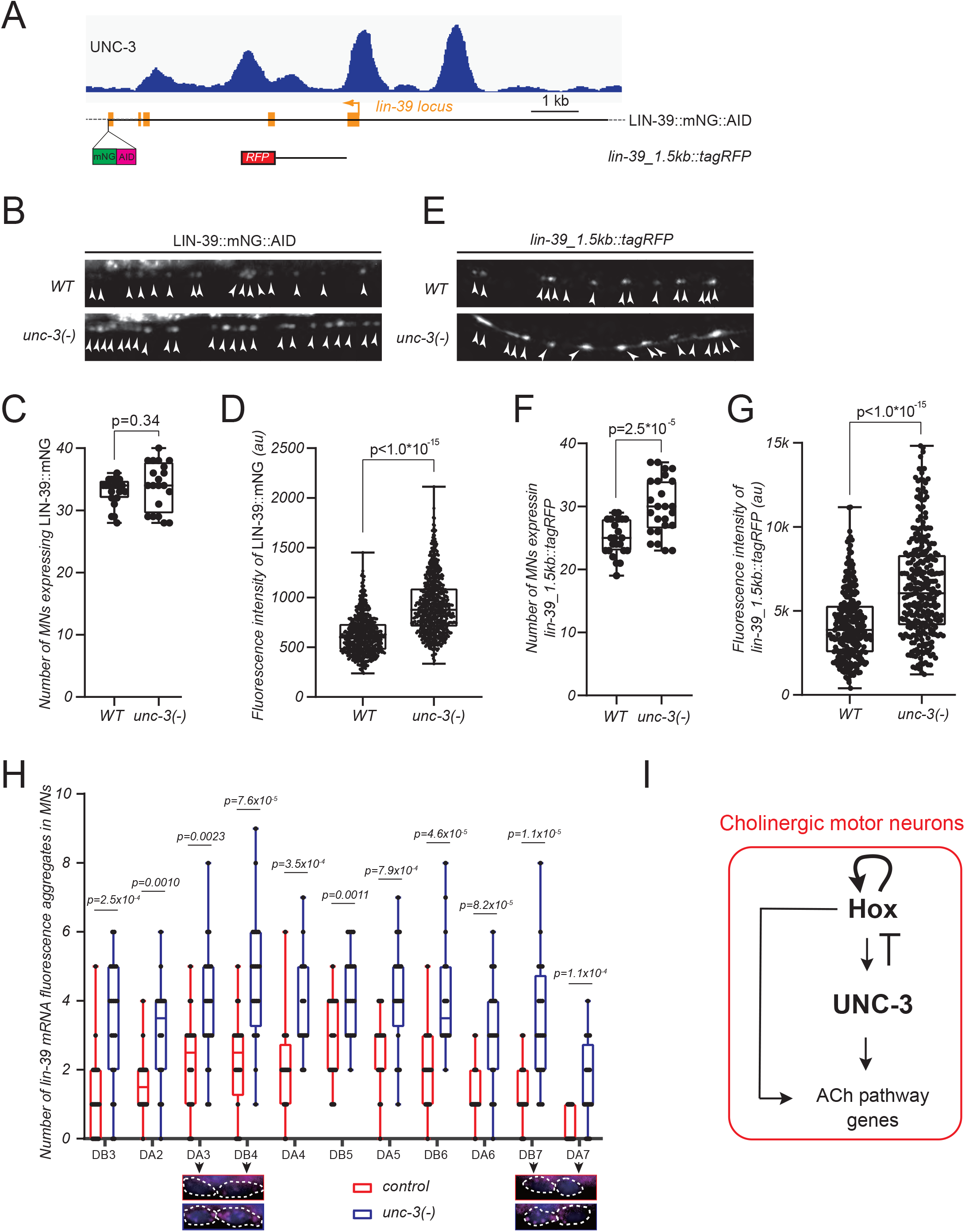
UNC-3 (Collier/Ebf) prevents high levels of Hox gene (*lin-39, mab-5*) expression. **A**. Genetic locus of *lin-39* overlay with ChIP-seq track of UNC-3. Molecular nature of *lin-39* reporter gene LIN-39::mNG::AID, and *lin-39_1*.*5kb::tagRFP* is annotated overlying below *lin-39* locus. **B-D**. Representative images (**B**) and quantifications of the number of MNs expressing LIN-39::mNG::AID (**C**) and the fluorescence intensity (**D**) in *WT* versus *unc-3(-)* background. **E-G**. Representative images (**E**) and quantifications of the number of MNs expressing *lin-39_1*.*5kb::tagRFP* (**F**) and the fluorescence intensity (**G**) in *WT* versus *unc-3(-)* background. For images in panel **B** and **E**, cell body positions are pointed out by white arrowheads. For quantifications, box and whisker plots were used with presentation of all data points. Unpaired t-test with Welch’s correction was performed and p-values were annotated. In panel **C** and **F**, N>20. In panel **D** and **G**, single neuron expressing reporter gene is scored for fluorescence intensity and plotted as individual data point in graph. N>600. See **Material and Methods** for details on fluorescence intensity quantification. **H**. Quantification of the number of *lin-39* mRNA fluorescence aggregates in individual cholinergic MNs via single molecule RNA Fluorescence in situ hybridization (smRNA-FISH) in WT versus *unc-3(-)* background with representative zooming-in images of DA3, DB4, DB7, and DA7. WT data is colored red versus mutant in blue for quantification. Box and whisker plots were used with presentation of all data points. Unpaired t-test with Welch’s correction was performed and p-values were annotated. N>20. Images are shown in merged channels of *lin-39* mRNA probe stained in magenta with DAPI counter stained the nucleus.

Taken together, our findings uncover an intricate gene regulatory network for maintenance of cholinergic identity in *C. elegans* motor neurons. This network contains a positive feedforward loop - Hox proteins (LIN-39 and MAB-5) not only directly activate the expression of ACh pathway genes, but also control the expression of *unc-3*, a known regulator of cholinergic identity in motor neurons (**Fig. 7I**). Moreover, Hox proteins LIN-39 and MAB-5 maintain their expression in motor neurons through transcriptional autoregulation, which is counterbalanced by UNC-3 (negative feedback), thereby generating a homeostatic control mechanism that ensures optimal levels of Hox gene expression (**Fig. 7I)**

## DISCUSSION

Hox genes play fundamental roles in early patterning and body plan formation, but their functions in later stages of development and post-embryonic life remain largely unknown. In the context of the nervous system, Hox genes are known to control early events, such as specification, survival, and/or migration of progenitor cells and young post-mitotic neurons [17-21]. Here, we identify a new role for Hox proteins during later stages of nervous system development and post-embryonic life, extending their functional repertoire beyond early patterning. Using *C. elegans* nerve cord motor neurons as a model, we found that Hox gene activity is continuously required, from development through adulthood, for the control of NT identity - a key function that ensures communication of neurons with their post-synaptic targets. Moreover, Hox genes operate in a positive feedforward loop to safeguard cholinergic motor neuron identity. That is, they not only control directly ACh pathway genes (e.g., *unc-17/VAChT, cho-1/ChT*), but also the expression of *unc-3* (*Collier/Ebf*), a master regulator of motor neuron identity [12]. Lastly, we propose a homeostatic mechanism that maintains Hox gene expression at optimal levels over time, thereby ensuring robust expression of ACh pathway genes in motor neurons.

### Hox genes *lin-39* (*Scr/Dfd, Hox4-5*) and *mab-5 (Antp, Hox6-8)* can function as terminal selectors of cholinergic identity in motor neurons

During the last steps of nervous system development, individual neuron types acquire their NT identity, and perhaps most important, such identity must be maintained throughout post-embryonic life. Transcription factors that are continuously expressed in individual neuron types and act directly to co-regulate the expression of NT pathway genes have been termed “terminal selectors” [40-42]. Terminal selectors have been described thus far for various neuron types in *C. elegans*, flies, simple chordates and mice, indicating their deep evolutionary conservation [43, 44]. Their sustained expression in specific neuron types suggests they are continuously required to maintain the expression of NT pathway genes. However, such continuous requirement has been experimentally demonstrated only for a handful of terminal selectors to date [45].

Recent work in *C. elegans*, flies and mice indicates that certain Hox genes are continuously expressed, from development through adulthood, in distinct neuron types [22-28]. Functional studies in peptidergic neurons in *Drosophila*, as well as hindbrain and spinal neurons in mice, showed that Hox proteins control early facets of neuronal development (e.g., specification, survival, migration, connectivity) [17-21, 46], but whether they function as terminal selectors remains unknown - in part due to a lack of temporally controlled studies for Hox gene inactivation later in life. Through constitutive (genetic null alleles) and post-embryonic (inducible protein depletion) approaches, we found that the Hox gene *lin-39* is continuously required to maintain the cholinergic identity of *C. elegans* motor neurons. Biochemical (ChIP-Seq) and genetic evidence strongly suggests that LIN-39 and another Hox protein (MAB-5) act directly to co-regulate the expression of multiple genes (*unc-17/VAChT, cho-1/ChT, ace-2/AChE*), whose protein products constitute integral components of the ACh biosynthetic pathway. Collectively, these findings indicate that Hox genes can function as terminal selectors of cholinergic motor neuron identity in the *C. elegans* nerve cord, thereby uncovering a new role for Hox during post-embryonic life.

Because Hox genes are expressed in various neuron types in *C. elegans* [28], we surmise they may function as terminal selectors in other neurons as well. Supporting this scenario, expression of a single marker of serotonergic identity (*tph-1*, tryptophan hydroxylase 1 [TPH1]) is affected in CP neurons of *lin-39* mutant animals [47, 48]. Similarly, expression of a dopaminergic identity marker (*cat-2*/tyrosine hydroxylase [TH]) is affected in tail sensory neurons of animals lacking activity of the posterior Hox gene *egl-5* (*AbdB*) [49]. Additional work is needed, however, to determine whether (a) LIN-39 and EGL-5 are continuously required in those neurons, and (b) they act directly to regulate additional genes involved in serotonin and dopamine biosynthesis, respectively.

### Hox proteins extend the list of homeodomain proteins acting as terminal selectors

Homeodomain proteins are defined by a 60 amino acid motif (homeodomain) that directly contacts DNA [50]. Based on sequence similarities and/or the presence of other domains, several subfamilies (e.g., HOX, LIM, POU, PRD) of homeodomain proteins have been identified in every animal genome. In *C. elegans*, a comprehensive expression analysis of all its 102 homeodomain proteins revealed that unique combinations of homeodomain proteins are found in all neuron types of the worm [51]. Accumulating evidence in *C. elegans* suggests that members of LIM, POU, and PRD subfamilies can function as terminal selectors in specific neuron types [52]. Here, we show that members of the HOX subfamily act as terminal selectors in cholinergic motor neurons. Subsequently, an overarching principle emerges, i.e., a single transcription factor family, the homeodomain proteins, is broadly used in the *C. elegans* nervous system to control neuronal identity by acting as terminal selectors. Supporting the evolutionary conservation of this theme, a number of LIM and POU homeodomain transcription factors can function as terminal selectors in the mouse nervous system [6, 7, 53-55]. Future studies will determine whether Hox proteins act as terminal selectors in vertebrate neurons.

### Hox proteins cooperate with the terminal selector UNC-3 (Collier/Ebf) to control motor neuron cholinergic identity

Besides NT identity, terminal selectors are known to control additional, neuron type-specific features of terminal differentiation, such as expression of ion channels, neuropeptides, NT receptors, cell adhesion molecules [40, 42]. We find this to be the case for LIN-39. In sex-specific motor neurons that control egg laying, LIN-39 is required for the expression ACh pathway genes and additional terminal differentiation markers (e.g., *glr-5/GluR, ida-1/PTPRN*). Similarly, in motor neurons necessary for locomotion, LIN-39 controls ACh pathway genes (this study) and a handful of terminal differentiation markers (e.g., *del-1*/sodium channel SCNN1, *slo-2*/potassium-sodium activated channel KCNT) [25, 31]. But how can the same Hox protein (LIN-39) operate as a terminal selector in two different types of motor neurons?

Our findings support the idea of distinct combinations of transcription factors acting as terminal selectors in different neuron types. In motor neurons that control locomotion, LIN-39 cooperates with MAB-5 and UNC-3, the latter is known to function as a terminal selector in these neurons [12]. Mechanistically, LIN-39 and UNC-3 control ACh pathway genes by recognizing distinct binding sites (motifs) on the genome. However, whether Hox and UNC-3 are being recruited to these sites in an additive or cooperative manner remains unresolved. In VC motor neurons that control egg-laying, UNC-3 is not expressed [12]. However, HLH-3 (bHLH transcription factor of the Achaete-Scute family) and LIN-11 (LIM homeodomain) are selectively expressed in VC neurons [56, 57], constituting putative terminal selectors that can potentially cooperate with LIN-39 to control VC cholinergic identity.

### Hox and UNC-3 in a positive feed-forward loop (FFL): a form of redundancy engineering to ensure robust expression of cholinergic identity genes

Complex gene regulatory networks are composed of simple gene circuits called “network motifs”[36]. One of the most widely used motifs is the feed-forward loop (FFL), where transcription factor X activates a second transcription factor Y, and both activate their shared target gene Z (**Fig. 5K**). Our findings uncovered a FFL in cholinergic motor neurons, where a Hox protein (LIN-39) activates UNC-3, and both activate their shared targets (Z = ACh pathway genes) (**Fig. 5K**). Although FFLs have been described across phylogeny, their biological functions to date have been experimentally tested primarily in bacteria [36-38]. Based on systems biology classifications, there are 8 different types of FFLs – each characterized by the signs (“+” for activation, “–” for repression) of transcriptional interactions within the motif. The FFL in motor neurons is coherent type 1 because all interactions are activating (**Fig. 5K**).

Why is there a need for this type of FFL in *C. elegans* motor neurons? Computational models and studies in bacteria indicate that a coherent type 1 FFL can ensure robust gene expression against perturbations through a process called “sign-sensitive delay” [38, 58, 59]. That is, a perturbation can decrease the levels of transcription factor X (LIN-39), but Z (ACh pathway genes) responds only at a delay once X levels decrease. The delay is due to the presence of Y (UNC-3). After X is decreased, it takes time for Y levels to decrease (depending on Y’s degradation rate) to a level insufficient to activate Z. Hence, the coherent type 1 FFL can be viewed as a form of redundancy engineering (or filter) to protect the shared target genes (Z) from fluctuations in the input (transcription factor X). It makes the system insensitive to brief periods during which X (LIN-39) is decreased. We therefore speculate that LIN-39 (X) and UNC-3 (Y) operate in a coherent type 1 FFL to maintain robust expression levels of ACh pathway genes in motor neurons throughout life (**Fig. 5K**).

This FFL forms the backbone of the transcriptional mechanism we identified in cholinergic motor neurons, but it does not function in isolation. Rather, it is embedded within at least two additional loops: (a) a positive feedback loop that activates LIN-39 (X) expression through transcriptional autoregulation, and (b) negative feedback provided by UNC-3 (Y) to reduce LIN-39 expression. The significance of both is discussed below.

### A two-component design principle for homeostatic control of Hox gene expression in motor neurons

In the context of early development, Polycomb Group (PcG) and trithorax Group (trxG) genes determine the levels and spatial expression of Hox genes with PcG proteins acting as repressors and trxG as activators [60-64]. In early nervous system development, Hox gene expression is initially established through morphogenetic gradients, and further refined via cross-regulatory Hox interactions [21, 65]. However, the mechanisms that maintain Hox gene expression in post-mitotic neurons during later developmental and post-embryonic stages remain unknown.

Our constitutive (genetic null alleles) and temporally controlled (auxin-inducible protein depletion) approaches combined with CRISPR/Cas9-mediated mutagenesis of Hox binding sites in the *lin-39* locus strongly suggest that LIN-39 maintains its own expression in motor neurons through transcriptional autoregulation (**Fig. 6**). Subsequently, LIN-39 is able to maintain the expression of ACh pathway genes by operating at the top of the FFL discussed in the previous section. However, transcriptional autoregulation can result - over time - in accumulation of high levels of LIN-39, possibly leading to detrimental effects in motor neurons. Indeed, LIN-39 overexpression specifically in motor neurons caused “mixed identity”, i.e., genes normally expressed in GABA neurons became ectopically expressed in motor neurons [31]. Hence, LIN-39’s transcriptional autoregulation must be counterbalanced through a mechanism that limits its production. We find that the terminal selector UNC-3 prevents high levels of *lin-39* expression in motor neurons, thereby counterbalancing LIN-39’s transcriptional autoregulation and avoiding its perpetual accumulation. From the perspective of Hox binding site affinity [66, 67], this mechanism may enable LIN-39 (when present at optimal levels) to bind to high-affinity sites in the *cis*-regulatory region of ACh pathway genes. The negative feedback by UNC-3 that prevents high levels of LIN-39 is potentially needed to avoid LIN-39’s binding to low-affinity sites in the *cis*-regulatory region of alternative (e.g., GABA) identity genes.

Altogether, we identified a two-component design principle (positive feedback of LIN-39 to itself, negative feedback from the terminal selector UNC-3) for the homeostatic control of Hox gene expression in MNs. These two components are embedded in a FFL, which we propose to be necessary for maintenance of NT identity. Because Hox genes are continuously expressed in the adult fly, mouse, and human nervous systems, the gene regulatory mechanism described here for the control of NT identity may be broadly applicable.

## MATERIALS AND METHODS

### *C. elegans* strains

Worms were grown at 15°C, 20°C or 25°C on nematode growth media (NGM) plates seeded with bacteria (*E*.*coli* OP50) as food source (Brenner, 1974). Mutant alleles used in this study: *unc-3 (n3435) X, lin-39 (n1760) III, mab-5 (e1239) III, ced-3 (n1286) IV*. CRISPR-generated alleles: *unc-3 (ot837 [unc-3::mNG::AID]) X, lin-39 (kas9 [lin-39::mNG::AID]) III*. All reporter strains used in this study are shown in **Supplementary Table 1**.

### Generation of transgenic reporter animals

Reporter gene fusions for *cis*-regulatory analysis of terminal identity genes were made using either PCR fusion (Hobert, 2002) or Gibson Assembly Cloning Kit (NEB #5510S). Targeted DNA fragments were fused (ligated) to *tagrfp* coding sequence, which was followed by *unc-54 3’ UTR*. Mutations or Deletions targeting LIN-39 binding sites were introduced via mutagenesis PCR. The product DNA fragments were either injected into young adult *pha-1(e2123)* hermaphrodites at 50ng/µl using *pha-1* (pBX plasmid) as co-injection marker (50 ng/µl) and further selected for survival, or injected into young adult N2 hermaphrodites at 50ng/µl (plus 50ng/µl pBX plasmid) using *myo-2::gfp* as co-injection marker (3 ng/µl) and further selected for GFP signal.

### Targeted genome engineering

To generated *776bp_HOX_MUT::mNG::AID* allele, CRISPR/Cas9 genome editing was employed to introduce targeted mutation to the *kas9 allele* at *lin-39* gene locus. Editing was performed by SunyBiotech.

### Temporally controlled protein degradation

AID-tagged proteins are conditionally degraded when exposed to auxin in the presence of TIR1 (Zhang et al., 2015). Animals carrying auxin-inducible alleles of *lin-39 (kas9 [lin-39::mNG::AID])* were crossed with *ieSi57* that expresses TIR1 ubiquitously or with otTi28 that expresses TIR-1 pan-neuronally. Auxin (indole-3-acetic acid [IAA]) was dissolved in ethanol (EtOH) to prepare 400 mM stock solutions which were stored at 4°C for up to one month. NGM agar plates were poured with auxin or ethanol added to a final concentration of 4 mM and allowed to dry overnight at room temperature. OP50 was cultured for the following days before transferring worms onto the plates. To induce protein degradation, worms of the experimental strains were transferred onto auxin-coated plates and kept at 25°C. As a control, worms were transferred onto EtOH-coated plates instead. Auxin solutions, auxin-coated plates, and experimental plates were shielded from light.

### Single Molecule RNA in situ Hybridization (sm RNA-FISH)

Egg prep was performed before harvesting the synchronized L1 worms. Following that, the worms were fixed with 4% paraformaldehyde (PFA), washed twice with PBS, and permeabilized with 70% ethanol at 4°C overnight. Then, samples are proceeded to hybridization with diluted mRNA probes purchased through Stellaris (unc-3: unc-3 best Quasar 670 (SMF-1065-5); lin-39: lin-39 best Quasar 670 (SMF-1065-5)) in purchased hybridization buffer (Stellaris® RNA FISH Hybridization Buffer (SMF-HB1-10)) for desired time following manufacturing protocols. Then, sample was stained with DAPI, washed, and equilibrated with GLOX buffer. And finally, worms were imaged at fluorescence microscope (Zeiss, Axio Imager.Z2).

### Microscopy

Worms were anesthetized using 100mM of sodium azide (NaN3) and mounted on a 4% agarose pad on glass slides. Images were taken using an automated fluorescence microscope (Zeiss, Axio Imager.Z2).Acquisuisition of Several Z-Stack Images(each ∼1 µm thick) was taken with Zeiss Axiocam 503 mono using the ZEN software (Version 2.3.69.1000, Blue edition). Representative images are shown following max-projection of 1-8 µm Z-stacks using the maximum intensity projection type. Image reconstruction was performed using Image J software (Schindelin et al., 2012).

### Motor neuron identification

Motor neuron (MN) subtypes were identified based on combinations of the following factors: (i) co-localization with fluorescent markers, with Known expression Pattern, (ii) invariant cell body position along the ventral nerve cord, or relative to other MN subtypes, (iii) MN birth order, and (iv) number of MNs that belong to each subtype.

### Bioinformatic analysis

To predict the UNC-3 binding site (COE motif) in the *cis*-regulatory region, we used the MatInspector program from Genomatix (Cartharius et al., 2005). The Position Weight Matrix (PWM) for the LIN-39 binding site is catalogued in the CIS-BP (Catalog of Inferred Sequence Binding Preferences database) (Weirauch et al., 2014). To identify putative LIN-39 sites on the *cis*-regulatory regions, we used FIMO (Find Individual Motif Occurrences)(Grant et al., 2011), which is one of the motif-based sequence analysis tools of the MEME (Multiple Expectation maximization for Motif Elicitation) bioinformatics suite (http://meme-suite.org/). The p-value threshold for the analysis was set at p<0.001 *for all genes except for lin-39 autoregulation sites where we set* p<0.01.

### Fluorescence Intensity (FI) Quantification

To quantify FI of individual MNs in the VNC, images of worms from different genetic background were taken with same imaging parameters through relative complete z-stacks that cover entire range of cell body. Image stacks were then processed and quantified for FI via FIJI. The focal plane in Z-stack that has the brightest FI is selected for quantification to minimize background signals. Cell outline was manually selected, and FIJI will quantify the FI and area to get the mean value for FI. After quantifying the FI of all MNs of interest, representative average image background FI is quantified and subtracted from the individual MN FI quantifications to get the mean FI in arbitrary unit (a.u).

### Statistical analysis

For quantification, box and whisker plots were adopted to represent the quartiles in graph. The box includes data points from the first to the third quartile value with the horizontal line in box representing the mean value. Upper and lower limits indicate the max and min, respectively. Unpaired t-test with Welch’s correction was performed and p-values were annotated. Visulization of data and p-value calculation were perform via GraphPad Prism Version 9.2.0 (283)

## ACKNOWLEDGEMENTS

We thank the Caenorhabditis Genetics Center (CGC), which is funded by NIH Office of Research Infrastructure Programs (P40 OD010440), for providing strains. We are grateful to Oliver Hobert and members of the Kratsios lab for comments on this manuscript. This work was funded by two NIH grants to P.K (R01 NS116365-01, R01 NS118078-01).

